# Associations of Minor Histocompatibility Antigens with Clinical Outcomes Following Allogeneic Hematopoietic Cell Transplantation

**DOI:** 10.1101/2022.08.31.506092

**Authors:** Othmane Jadi, Hancong Tang, Kelly Olsen, Steven Vensko, Qianqian Zhu, Yiwen Wang, Christopher A Haiman, Loreall Pooler, Xin Sheng, Guy Brock, Amy Webb, Marcelo C. Pasquini, Philip L McCarthy, Stephen R. Spellman, Theresa Hahn, Benjamin Vincent, Paul Armistead, Lara E. Sucheston-Campbell

**Affiliations:** Lineberger Comprehensive Cancer Center, University of North Carolina at Chapel Hill, CB# 7295, Chapel Hill, NC, 27599-7295, USA; College of Pharmacy, The Ohio State University, Columbus Ohio, USA; Department of Biostatistics and Bioinformatics, Roswell Park Comprehensive Cancer Center, Buffalo, NY, USA; Quantitative Sciences Unit, Department of Medicine, Stanford University, Palo Alto, CA, 34304, USA; Department of Preventive Medicine, University of Southern California; The Center for Genetic Epidemiology, USC, Los Angeles, CA 90033; Department of Biomedical Informatics, The Ohio State University, Columbus, OH 43210; Center for International Blood and Marrow Transplant Research, Medical College of Wisconsin, Milwaukee, WI; Center for International Blood and Marrow Transplant Research, National Marrow Donor Program, Minneapolis, MN; Department of Cancer Prevention & Control, Roswell Park Comprehensive Cancer Center, Buffalo, NY, USA; Division of Hematology, Department of Medicine, UNC School of Medicine, Chapel Hill, NC, USA; College of Veterinary Medicine, The Ohio State University, Columbus, Ohio, USA

## Abstract

The role of minor histocompatibility antigens (mHAs) in mediating graft versus leukemia (GvL) and graft versus host disease (GvHD) following allogeneic hematopoietic cell transplantation (alloHCT) is recognized but not well-characterized. By implementing improved methods for mHA prediction in two large patient cohorts, this study aimed to comprehensively explore the role of mHAs in alloHCT by analyzing whether (1) the number of predicted mHAs, or (2) individual mHAs are associated with clinical outcomes using multi-variate survival models corrected for multiple testing. Cox proportional hazard results showed that patients with a class I mHA count greater than the population median had an increased hazard of GvHD mortality (HR=1.39, 95%CI 1.01, 1.77, P=0.046). Competing risk analyses identified the class I mHAs DLRCKYISL (gene *GSTP*), WEHGPTSLL (*CRISPLD2*) and STSPTTNVL (*SERPINF2*) were associated with increased GVHD death (HR=2.84, 95%CI 1.52, 5.31, P=0.01), decreased leukemia-free survival (LFS) (HR=1.94,95%CI 1.27, 2.95, P=0.044), and increased disease-related mortality (DRM) (HR=2.32, 95%CI 1.5, 3.6, P=0.008), respectively. One class II mHA YQEIAAIPSAGRERQ (T*ACC2*) was associated with increased risk of treatment-related mortality (TRM) (HR=3.05, 95%CI 1.75, 5.31, P=0.02). WEHGPTSLL and STSPTTNVL were present in conjunction within HLA haplotype B*40:01-C*03:04 and showed a positive dose-response relationship with increased all-cause mortality and DRM and decreased LFS, indicating these two mHAs contribute to risk of mortality in an additive manner. Our study reports the first large scale investigation of the associations of predicted class I and class II mHA peptides with clinical outcomes following alloHCT.

**Graphical Abstract:** 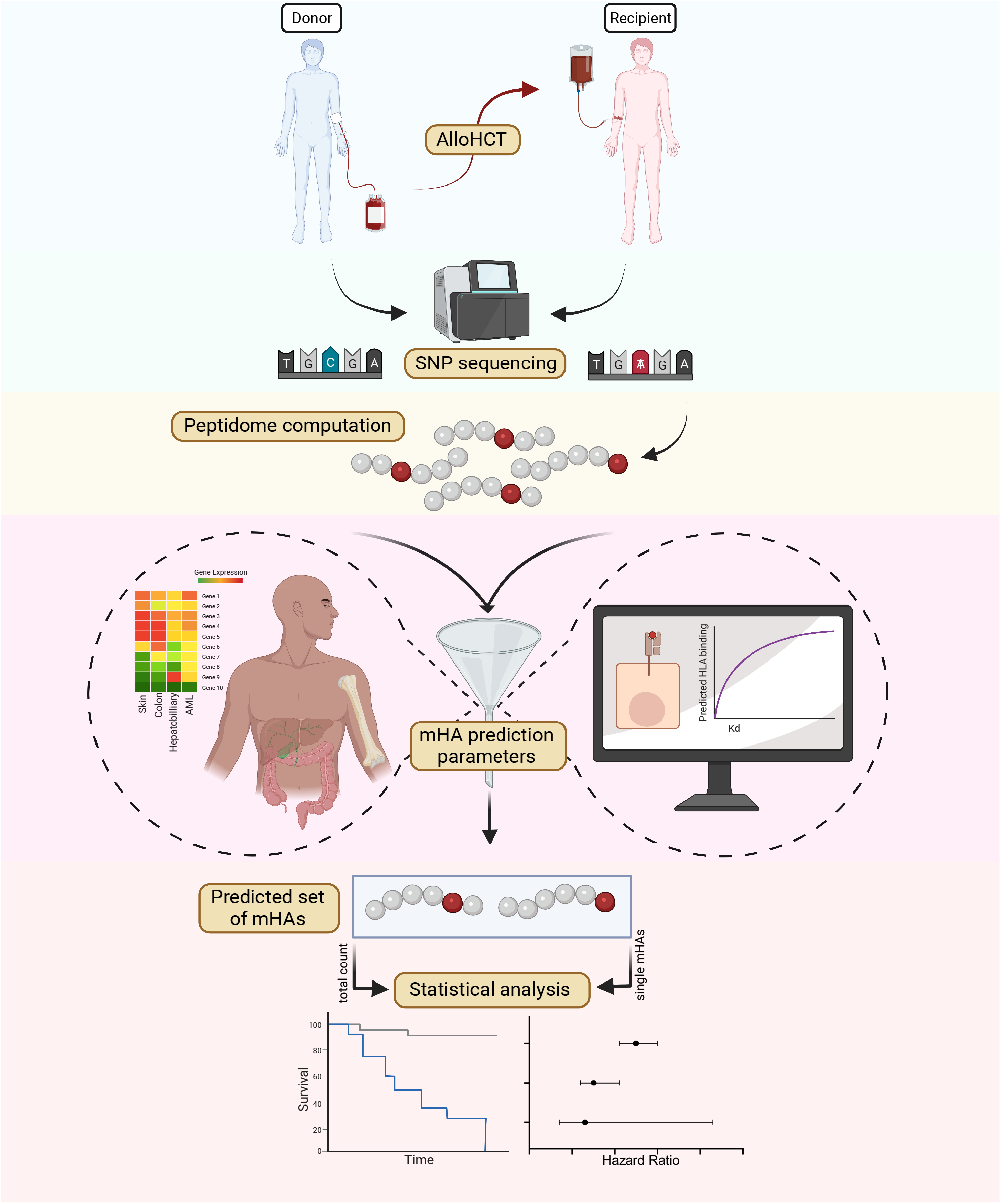

## INTRODUCTION

Allogeneic hematopoietic cell transplantation (alloHCT) involves the infusion of donor hematopoietic progenitor cells and T lymphocytes into transplant recipients. Successful treatment of hematologic neoplasms with alloHCT depends on donor T cell responses for eradication of tumor cells and relapse prevention.^1^ This therapeutic graft versus leukemia (GvL) effect is driven by the recognition of tumor cells as foreign by engrafted T cells. Minor histocompatibility antigens (mHAs) presented on tumor cells are targeted by donor T cells and are thought to be central mediators of GvL in HLA-matched transplants.^2,3^ mHAs are immunogenic peptides derived from differences in genetic variation, usually single nucleotide polymorphisms (SNPs), between an alloHCT donor and patient (recipient).^4^ T-cell receptors recognize cognate mHAs presented by major histocompatibility complexes (MHCs) on recipient cancer cells, which activates donor T-cells to kill these malignant cells. In contrast, graft versus host disease (GvHD) develops when donor T-cells react to mHAs presented on recipient normal cells. Because T-cell responses are dependent on peptide-MHC (pMHC) complexes, the immunogenicity of an mHA depends on both the peptide itself and the MHC (i.e., HLA) allele presenting it.

Increased disease relapse and decreased GvHD in identical-twin transplants compared to HLA-identical sibling transplants demonstrated the potential role of mHAs in alloHCT outcomes due to the assumed paucity of mHAs in identical-twin transplants.^3,5^ However, it is unclear whether GvL and GvHD are predominantly mediated by the cumulative effect of many mHAs and/or individual immunodominant mHAs. Several mHAs have been identified through isolation of antigen-specific T cells from alloHCT recipients, peptide elution from HLA molecules, or cDNA library screening.^4^ However much of the focus has been on isolating potential immunotherapeutic targets rather than identifying mHAs that drive clinical outcomes following alloHCT. Prior studies have identified SNPs associated with outcomes, but these studies identified genomic variants rather than mHA peptides.^6,7^

More recently, “reverse immunology” approaches have utilized genomic sequencing & other bioinformatic tools to predict the entire mHA landscape of donor-recipient pairs (DRPs) following alloHCT.^8-11^ This approach has allowed studies to assess the impact of the predicted mHA repertoire on outcomes. A small cohort analysis found no association between the number of predicted mHAs and clinical outcomes.^9^ Another study found no association between predicted HLA binding efficiency, used as a proxy for mHA quantification, and clinical outcomes.^12^ One study did find a significant association between SNP mismatching and acute GVHD in matched-related DRPs.^13^ However, these studies have major limitations due to small patient cohorts or imprecise predictions of mHA. No prior study has predicted mHAs in a large patient cohort to investigate their association with clinical outcomes after alloHCT.

Efforts to translate donor-recipient genetic differences into putative mHAs are also challenged by the small fraction of SNPs that makes up the mHA repertoire (0.5%) and the small fraction of possible n-mer peptides that binds HLA molecules (<0.1%).^11,14^ Candidate mHAs have previously been identified using only predicted HLA binding affinity (e.g. <500nM), but this correlates weakly with actual immunogenicity.^15,16^ Optimized methods for the prediction of tumor neoantigens have the potential to also improve predictions of immunogenic mHAs^16^. A recent study established a set of parameters to optimize the prediction of therapeutically relevant neoantigens based on a set of peptide features.^17^ Applying this model to the prediction of mHAs may allow us to better analyze the role of mHAs in alloHCT by improving predictions of clinically significant mHAs.

While prior analyses have identified common SNPs and rare variants associated with post-alloHCT outcomes, these were HLA-agnostic analyses and focused on variants rather than peptides.^6,7^ No previous study has examined the association between mHA peptides and clinical outcomes in a large patient cohort. Here, we hypothesized that an increased mHA count in a DRP and individual mHAs would associate with increased GvHD mortality and/or lower disease related mortality due to decreased relapse. We assessed this by analyzing the existing Determining the Influence of Susceptibility COnveying Variants Related to one-Year mortality after BMT (DISCOVeRY-BMT) dataset with two goals: (1) to determine whether the number of predicted mHAs correlate with clinical outcomes; (2) to identify novel individual mHAs that are significantly associated with clinical outcomes through HLA-specific analyses.

## METHODS

### Clinical Data Source

The study population consisted of DISCOVeRY-BMT cohorts (Table 1).^7,18^ Patients in both cohorts were treated for acute myeloid leukemia (AML), acute lymphoblastic leukemia (ALL), or myelodysplastic syndrome (MDS) with alloHCT from HLA-matched unrelated donors. Cohort 1 consists of 10/10 HLA matched (HLA-A, -B, -C, -DRB1, and -DQB1 loci) DRPs who underwent transplant from 2000 to 2008. Cohort 2 consists of 10/10 HLA matched DRPs from 2009 to 2011 and 8/8 HLA matched (HLA-A, -B, -C, and -DRB1 loci) DRPs from 2000 to 2011. Our study focused on patients with myeloid malignancies (AML and MDS), which comprised 78% of the DISCOVeRY-BMT dataset. Of our total population less than five percent of patients were of African, South Asian, East Asian, Pacific Islander or multiple ancestries, or Hispanic ethnicity; we present analysis results for only non-Hispanic European-American (EA) patients (n=2249) to reduce confounding effects from genetic ancestry. The 1-year clinical outcomes used in our analyses included overall survival, leukemia-free survival (LFS), treatment-related mortality (TRM), disease-related mortality (DRM), disease relapse, and mortality due to GvHD. GvHD mortality was defined as death driven or contributed to by GvHD, without evidence of primary disease progression or relapse. Survival outcomes were adjudicated by a consensus panel.^19^ All donors and recipients provided informed consent for use of their clinical data in research.

**Table 1.**
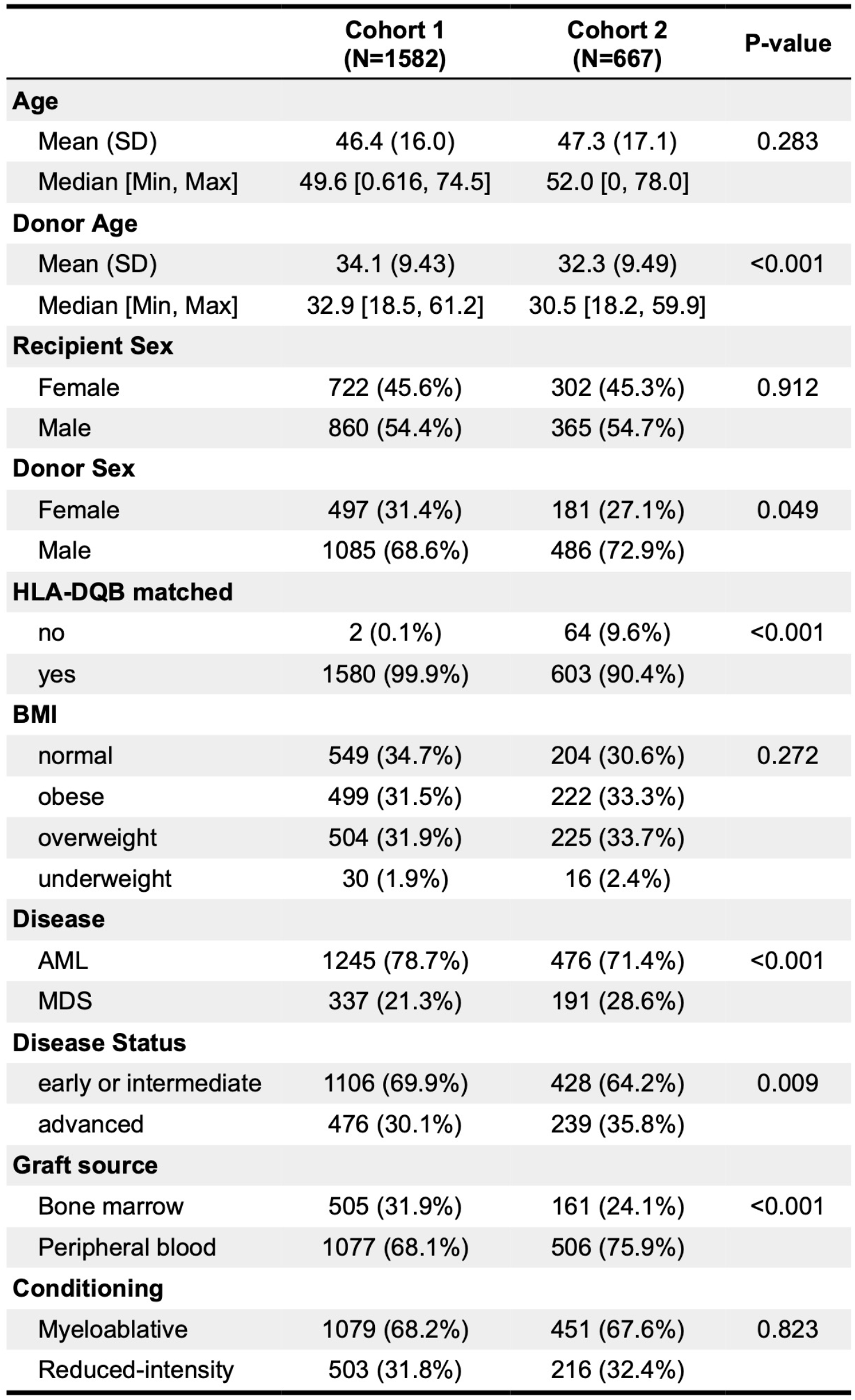
Patient characteristics for both study cohorts. P values for differences in cohort 1 and 2 were calculated using Pearson’s Chi-squared test.

### Genotyping, Imputation and Quality Control

Genotyping, imputation, and quality control have been previously described in detail.^19-23^ Briefly, DNA from blood samples of donors and recipients was genotyped using Illumina Human OmniExpress BeadChip containing ∼733,000 SNPs. Quality control (QC) was performed at both the variant and sample level. SNPs were removed if they had a missing call rate > 2%, a minor allele frequency < 0.001, or violated Hardy Weinberg Equilibrium proportions (p < 1 × 10^−4^). Sample-wise QC removed patients with a missing call rate > 2%, reported-genotyped sex mismatch, abnormal inbreeding coefficients, cryptic relatedness, or who were population outliers. SNPs were then imputed through the Sanger Imputation Service using the Haplotype Reference Consortium hg19/b37 reference genome.^24^ SNPs with minor allele frequency > 0.005 and imputation quality score > 0.9 were used for mHA predictions.

### Minor Histocompatibility Antigen Prediction

To predict mHAs in a DRP, we translated nonsynonymous SNP alleles present in alloHCT recipients but absent in their donor (mHA-mismatched SNPs) into all possible peptides of varying lengths that would encompass the altered amino acid. We computed peptides 8-11 amino acids long for class-I restricted mHAs, and 15-24 amino acids long for class-II restricted mHAs. Nonsynonymous SNPs were identified using ANNOVAR and translated into peptides using antigen.garnish.^25,26^ These peptides were then filtered based on source protein gene expression and recipient HLA binding affinity using thresholds established for tumor neoantigen prediction.^16,17^ Peptides were filtered for source protein expression of > 33 transcripts per million (TPM) in tissues most clinically relevant to alloHCT outcomes – skin, hepatobiliary, colon, and AML.^17^ RNAseq expression data were acquired from UCSC’s Xena Dataset repository.^27,28^ 8-11 nmers were filtered for HLA binding affinity < 34nM to any recipient HLA-A,-B,-C allele predicted by netMHCpan.^29^ Given differences in binding affinity hemodynamics between the class I and class II HLA, we relied on a related but different parameter to identify peptides most likely to be presented by class II HLA – the eluted ligand (EL) prediction score from netMHCIIpan.^30,31^ 15-24 nmers were filtered for percentile rank of EL prediction score < 2% to identify peptides strongly bound to HLA-DRB1.^30^ The final pool of 8-11 nmers were identified as class I mHAs and the 15-24 nmers as class II mHAs.

### Number of mHA association with survival outcomes

All statistical analyses were performed in R.^32^ Two-sample Kolmogorov-Smirnov test was used to test the distribution difference in the total mHA counts between the two cohorts.^32^ Survival models were used to investigate the association of 1-year clinical outcomes after transplant with mHA counts, both total number per DRP and membership in the top or bottom half of the mHA distribution. OS and LFS were analyzed using Kaplan-Meier and Cox proportional hazards (CoxPH) models with the R package Survival.^33^ TRM, DRM, relapse, and GvHD mortality were analyzed using Fine and Gray sub-distributional hazard models with the R package cmprsk to account for competing events.^34^ Model covariates were selected using bidirectional stepwise selection for each individual outcome.^7,21^ Spearman correlation was used to test correlations among model variables through the R package Hmisc.^35^ Cohorts 1 and 2 were analyzed independently with subsequent random-effects meta-analysis modeling of the two cohorts. Class I and class II mHAs were initially analyzed separately and subsequently together for outcomes with significant associations.

### Single and multi-mHA association analysis with survival outcomes

We stratified our individual mHA outcomes analysis by HLA type. To ensure adequate sample sizes for our analysis, we selected HLA alleles with a study population allele frequency >5% and filtered for mHAs with a >5% frequency among DRPs within each HLA allele. Each mHA was tested individually for association with clinical outcomes using the same models and covariates as above. We used Bonferroni correction to adjust significance levels based on the number of independent source SNPs and report the adjusted P values herein as P_B_. SNPs were considered independent if they had linkage disequilibrium (LD) value R^2^ < 0.2, calculated using the R package SNPRelate.^36^ If more than one mHA associated with clinical outcome within a HLA haplotype, we conducted a multi-mHA survival analysis. For every mHAs significantly associated with a clinical outcome, we analyzed whether the association was also observed in DRPs with the cognate HLA type in which both the donor and recipient contain the peptide (ie, the peptide was not characterized as a mHA).

### Data sharing statement

De-identified participant data are available at the Center for International Blood and Marrow Transplant (www.cibmtr.org). Both genotype and phenotype data will also be made available in dbGaP excluding 978 DRPs for which the informed consent is not compliant with the NIH Genomic Data Sharing Policy.

## RESULTS

### Patient characteristics and model covariates

Our study focused on non-Hispanic EA alloHCT recipients diagnosed with MDS or AML (n=2249, Table 1). Cohort 1 had a higher mean donor age, proportion of patients with AML, proportion of recipients receiving a bone marrow graft. Patients in cohort 2 were more likely to have advanced disease at the time of transplant and to be treated with a peripheral blood graft. There were no differences in OS between the two cohorts. Because they differed in various disease and treatment characteristics, cohorts 1 and 2 were analyzed independently with subsequent meta-analysis. The clinical covariates used in competing risk models showed associations with various outcomes consistent with previous analyses of these data (Supplemental Table 1).^18,21^

### Set of predicted mHAs

The distribution of class I mHA counts did not significantly differ between cohorts 1 and 2 (P=0.88), with medians of 40 and 39 mHAs, respectively (Figure 1A). The distribution of class II mHA counts did not significantly differ between cohorts 1 and 2 (P=0.51) (Figure 1B), with medians of 3049 and 3062 mHAs, respectively. The number of class I mHAs per DRP was not correlated with the number of mHA-mismatched SNPs between the donor and recipient (Figure 1C, Pearson’s r = 0.012, P=0.57). Across DRPs, the number of class II mHAs was weakly correlated with the number of mHA-mismatched SNPs (Figure 1D, Pearson’s r = 0.08, P<0.01). The numbers of class I and class II mHAs among DRPs were not correlated (Figure 1E, Pearson’s r = 0.013, P=0.53). The numbers of class I and II mHAs did not correlate with other clinical variables used in the survival models (Figure 1F).

**Figure 1.**
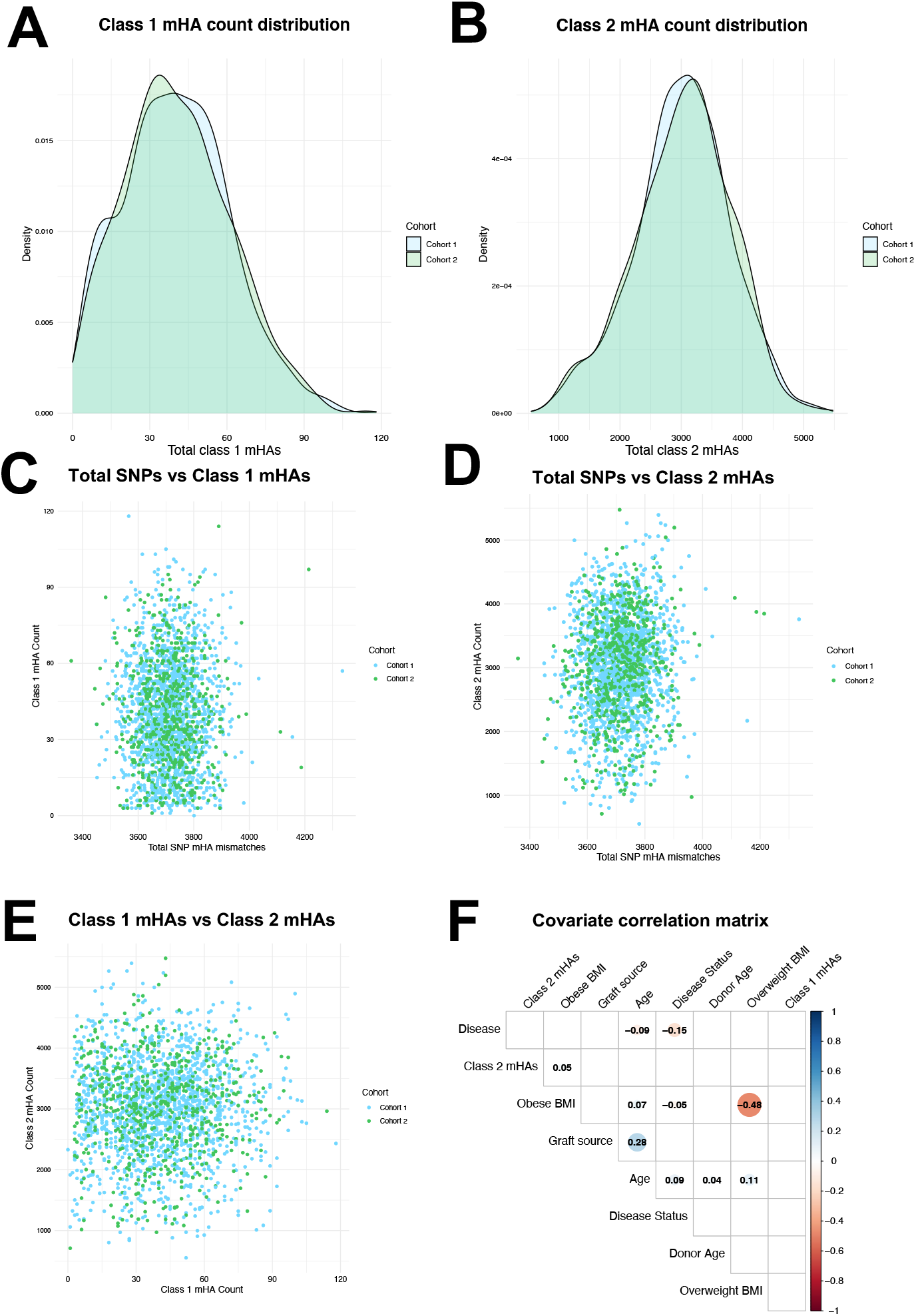
Distributions and correlations among predicted sets of mHAs. **(A)** Distribution of class 1 mHA counts is similar across both cohorts. **(B)** Distribution of class 2 mHA counts is similar across both cohorts. **(C)** There is no correlation between total between mHA-mismatched SNPs and the number of class 1 mHAs. **(D)** There is a weak correlation between total mHA-mismatched SNPs and the number of class 2 mHAs. **(E)** There is no correlation between the numbers of class 1 and class 2 mHAs. **(F)** There is no correlation between the numbers of mHAs and other covariates used in survival models. Correlations were calculated using Spearman’s Rho.

### Number of class I and class II mHAs associations

Patients with a class I mHA count greater than the population median had an estimated 39% increase in hazard of GvHD mortality compared to those with a count less than the median (HR=1.39, 95% CI = [1.01, 1.77], P=0.046) (Figure 2A). Class I mHA count quantile did not correlate with other clinical outcomes including overall survival, leukemia-free survival, relapse, DRM, and TRM. The number of class I mHAs as a quantitative variable did not correlate with any outcomes.

**Figure 2.**
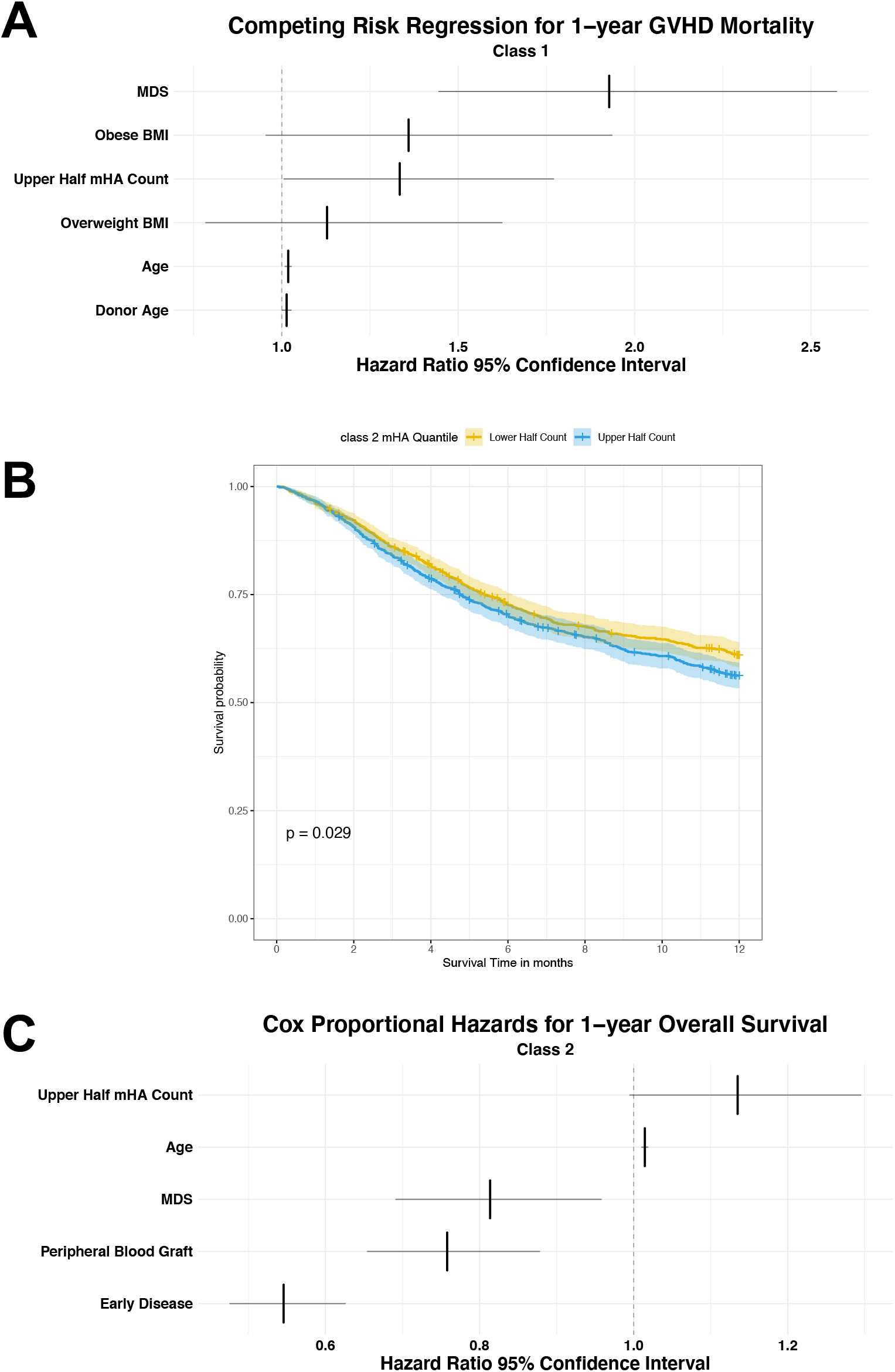
The number of mHAs is associated with various clinical outcomes. **(A)** Hazard ratios with 95% confidence intervals in a competing risk regression for 1-year mortality due to GvHD. Patients with a class 1 mHA count in the upper half of the population distribution had 39% increased hazard for 1-year GvHD mortality compared those with a count in the lower half (HR=1.39, 95% CI = [1.005, 1.772], P=0.046). **(B)** A Kaplan-Meier model shows that patients with a class 2 mHA count in the upper half had decreased overall survival compared to those with a count in the lower half (p=0.029). **(C)** Hazard ratios with 95% confidence intervals in a Cox-proportional hazards model show patients with a class 2 mHA count in the upper half had a 13% higher hazard for decreased overall survival (HR=1.135, 95% CI = [0.99, 1.30], P=0.06).

Patients with a class II mHA count greater than the population median had a decreased overall survival compared to those with a count less than the median (Figure 2B, Kaplan-Meier, p=0.029). However, this did not remain significant when adjusting for clinical covariates in a CoxPH model (HR=1.14, 95% CI = [0.99, 1.30], P=0.06, Figure 2C). Analysis of DRPs with class II mHA count in the upper quantile also showed evidence of decreased LFS (HR=1.12, 95% CI=[0.989,1.26], P=.07). Class II mHA count quantile did not correlate with other 1-year clinical outcomes. The number of class II mHAs as a quantitative variable did not correlate with outcomes.

A multivariate analysis including both class I and class II mHA counts together showed a similar but weaker association of class I mHA count quantile with increased GvHD mortality (HR=1.31, 95% CI = [0.983,1.754], P=0.065) and a consistent but not significant association of class II mHA count quantile with decreased overall survival (HR=1.13, 95% CI = [0.99, 1.29], P=0.06).

### Single mHA associations

We found 516 mHAs across 15 class I HLA alleles (HLA-A,-B,-C) that met frequency criteria for inclusion in the single mHA associations analysis. Two HLA alleles – A*01:01 and C*07:01 – had a small number of shared mHAs (>5% of DRPs), one and three respectively (Table 2).

**Table 2.**
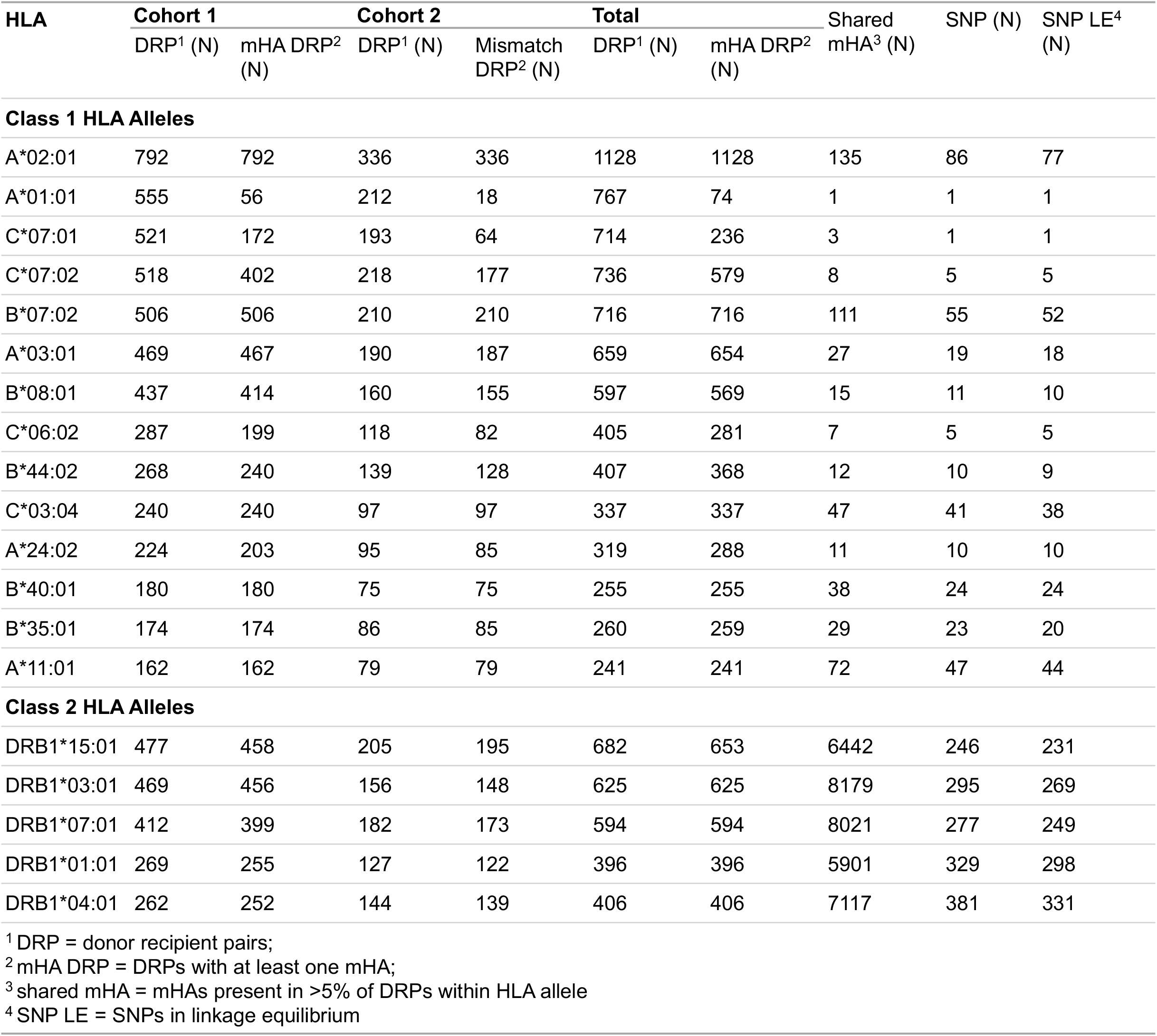
Number of DRPs, mHAs, and mHA-encoding SNPs within the most common HLA alleles. For class 1 and class II HLA alleles with a study population frequency >5%, we show the total number of DRPs and the number of DRPs with at least 1 mHA for each cohort and across both combined. For each HLA allele, we also show the total number of mHAs present in >5% DRPs (shared mHAs), shared mHA-encoding SNPs, and shared mHA-encoding SNPs in linkage disequilibrium.

| HLA | Cohort 1 |  | Cohort 2 |  | Total |  | Shared mHA<sup>3</sup> (N) | SNP (N) | SNP LE<sup>4</sup> (N) |
| --- | --- | --- | --- | --- | --- | --- | --- | --- | --- |
|  | DRP<sup>1</sup> (N) | mHA DRP<sup>2</sup> (N) | DRP<sup>1</sup> (N) | Mismatch DRP<sup>2</sup> (N) | DRP<sup>1</sup> (N) | mHA DRP<sup>2</sup> (N) |  |  |  |
| Class 1 HLA Alleles |  |  |  |  |  |  |  |  |  |
| A*02:01 | 792 | 792 | 336 | 336 | 1128 | 1128 | 135 | 86 | 77 |
| A*01:01 | 555 | 56 | 212 | 18 | 767 | 74 | 1 | 1 | 1 |
| C*07:01 | 521 | 172 | 193 | 64 | 714 | 236 | 3 | 1 | 1 |
| C*07:02 | 518 | 402 | 218 | 177 | 736 | 579 | 8 | 5 | 5 |
| B*07:02 | 506 | 506 | 210 | 210 | 716 | 716 | 111 | 55 | 52 |
| A*03:01 | 469 | 467 | 190 | 187 | 659 | 654 | 27 | 19 | 18 |
| B*08:01 | 437 | 414 | 160 | 155 | 597 | 569 | 15 | 11 | 10 |
| C*06:02 | 287 | 199 | 118 | 82 | 405 | 281 | 7 | 5 | 5 |
| B*44:02 | 268 | 240 | 139 | 128 | 407 | 368 | 12 | 10 | 9 |
| C*03:04 | 240 | 240 | 97 | 97 | 337 | 337 | 47 | 41 | 38 |
| A*24:02 | 224 | 203 | 95 | 85 | 319 | 288 | 11 | 10 | 10 |
| B*40:01 | 180 | 180 | 75 | 75 | 255 | 255 | 38 | 24 | 24 |
| B*35:01 | 174 | 174 | 86 | 85 | 260 | 259 | 29 | 23 | 20 |
| A*11:01 | 162 | 162 | 79 | 79 | 241 | 241 | 72 | 47 | 44 |
| Class 2 HLA Alleles |  |  |  |  |  |  |  |  |  |
| DRB1*15:01 | 477 | 458 | 205 | 195 | 682 | 653 | 6442 | 246 | 231 |
| DRB1*03:01 | 469 | 456 | 156 | 148 | 625 | 625 | 8179 | 295 | 269 |
| DRB1*07:01 | 412 | 399 | 182 | 173 | 594 | 594 | 8021 | 277 | 249 |
| DRB1*01:01 | 269 | 255 | 127 | 122 | 396 | 396 | 5901 | 329 | 298 |
| DRB1*04:01 | 262 | 252 | 144 | 139 | 406 | 406 | 7117 | 381 | 331 |
| <sup>1</sup> DRP = donor recipient pairs; |  |  |  |  |  |  |  |  |  |
| <sup>2</sup> mHA DRP = DRPs with at least one mHA; |  |  |  |  |  |  |  |  |  |
| <sup>3</sup> shared mHA = mHAs present in >5% of DRPs within HLA allele |  |  |  |  |  |  |  |  |  |
| <sup>4</sup> SNP LE = SNPs in linkage equilibrium |  |  |  |  |  |  |  |  |  |

Meta-analyses across both cohorts identified three class I mHAs and one class II mHA that were significantly associated with one-year clinical outcomes (Figures 3). In recipients with HLA-B*08:01, the mHA DLRCKYISL (*GSTP1*, SNP ID rs1695, ref. allele G) was associated with increased risk of GVHD mortality (HR=2.84, 95% CI = [1.52, 5.31], P_B_=0.01). Within HLA-B*40:01, WEHGPTSLL (*CRISPLD2*, rs12051468, ref. allele A) was associated with decreased LFS (HR=1.94, 95% CI = [1.27, 2.95], P_B_=0.044). Within HLA-C*03:04, STSPTTNVL (*SERPINF1*, rs1136287, ref. allele, C) was associated with increased risk of DRM (HR=2.32, 95% CI = [1.5, 3.6], P_B_=0.008). In recipients with HLA-DRB1*15:01, the class II mHA YQEIAAIPSAGRERQ (*TACC2*, rs11200385, alt. allele, A) was associated with increased risk of TRM (HR=3.05, 95% CI = [1.75, 5.31], P_B_=0.02). Importantly, these peptides were not associated with outcomes when present in both the donor and recipient of a DRP with the appropriate HLA allele (i.e., the recipient contained the peptide, but it did not meet our definition of a mHA).

**Figure 3.**
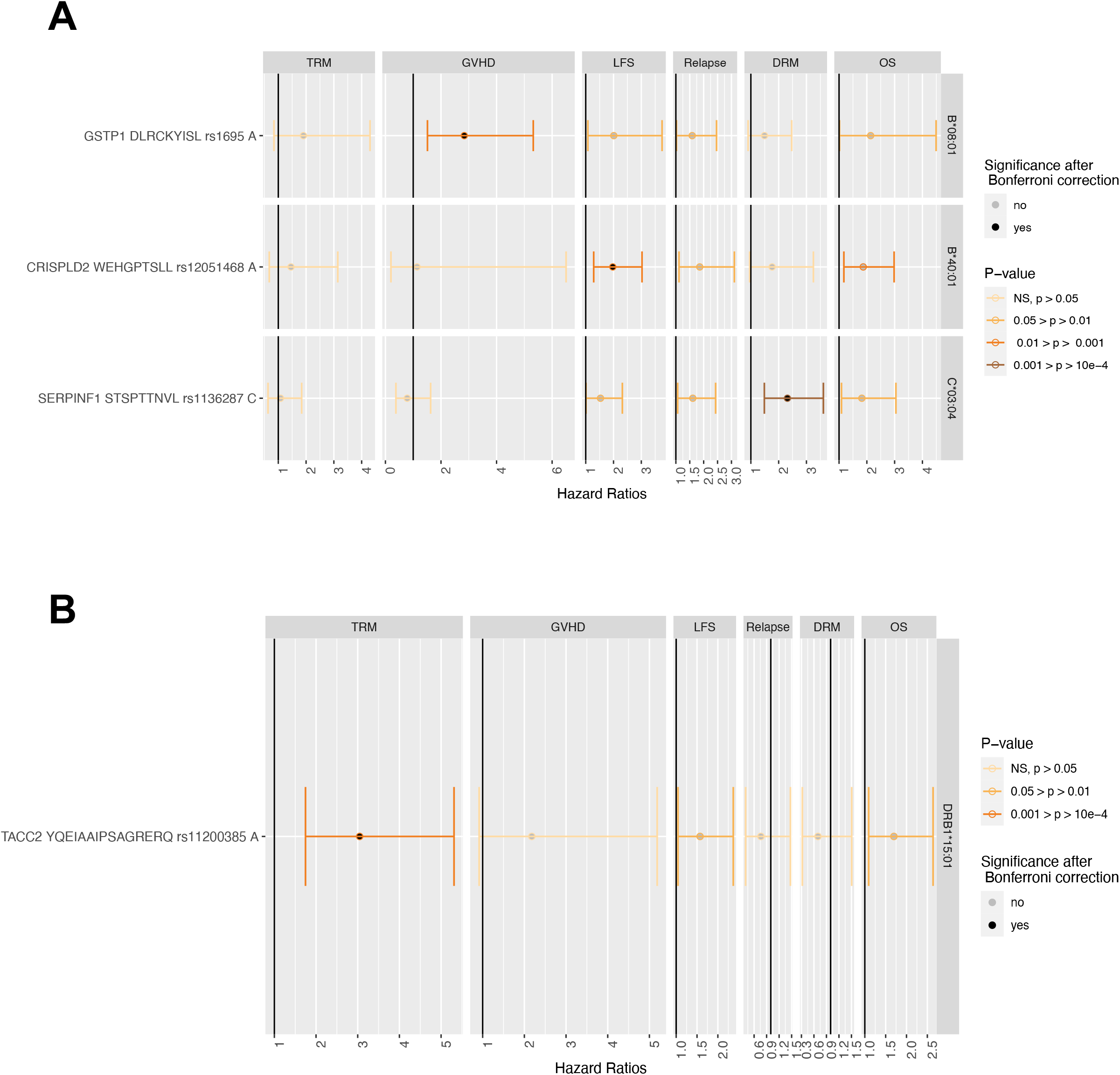
Individual class 1 and class 2 mHAs are associated with various clinical outcomes. For each mHA (row), we show hazard ratios with a 95% confidence interval in survival models of each clinical outcome (column) in a given HLA-allele. **(A)** mHA DLRCKYISL was associated with increased hazard of 1-year GvHD mortality in recipients with HLA B*08:01 (HR=2.84, 95% CI = [1.52, 5.31], P=1.1×10^−3^). WEHGPTSLL was associated with decreased leukemia free survival in patients with HLA B*40:01 (HR=1.94, 95% CI = [1.27, 2.95], P=2.016×10^−3^). STSPTTNVL was associated with increased risk of DRM in patients with HLA C*03:04 (HR=2.32, 95% CI = [1.5, 3.6], P=1.7×10^−4^). **(B)** YQEIAAIPSAGRERQ was associated with increased risk of TRM (HR=3.05, 95% CI = [1.75, 5.31], P=0.02) in patients with HLA DRB1*15:01.

The HLA alleles presenting WEHGPTSLL and STSPTTNVL, B*40:01 and C*03:04, make-up a haplotype present in 11% of EAs in the US.^37^ In our study population, 98% of DRPs with B*40:01 also have C*03:04, and 74% of DRPs with C*03:04 have B*40:01. We therefore analyzed the association of outcomes within this haplotype, comparing DRPs with both WEHGPTSLL and STSPTTNVL, or either mHA, to DRPs without these mHAs.

Among the 253 DRPs with HLA-B*40:01-C*03:04 (11% of DISCOVeRY-BMT cohorts), 14 DRPs had both mHAs, and 76 had either mHA (Supplemental Figure 1). Compared to having neither mHA (n=163), recipients with both mHAs had a significantly greater hazard of decreased 1-year overall survival (HR=3.58, 95% CI = [1.90, 6.76], P_B_=7.94×10^−5^), decreased 1-year leukemia free survival (HR=3.01, CI=[1.66,5.5], P_B_=2.96×10^−4^), increased DRM (HR = 3.65, 95% CI = [1.57, 8.51], P_B_= 0.003), and increased relapse (HR=2.85, 95% CI = [1.43, 5.68], P_B_=0.003) (Figures 4B-E, Supplemental Figure 2, Supplemental Table 2). Patients with only one mHA also had significantly greater hazard for these outcomes compared to patients with neither mHA, but to a lesser magnitude than patients with both mHAs. These patients had decreased 1-year overall survival (HR=1.69, 95% CI = [1.15, 2.49], P_B_=8×10^−3^), decreased 1-year leukemia free survival (HR=1.58, CI=[1.10, 2.3], P_B_=0.013), increased DRM (HR = 2.05, 95% CI = [1.21, 3.47], P_B_=0.008) and increased relapse (HR=1.71, 95% CI = [1.07, 2.74], P_B_=0.025).

**Figure 4.**
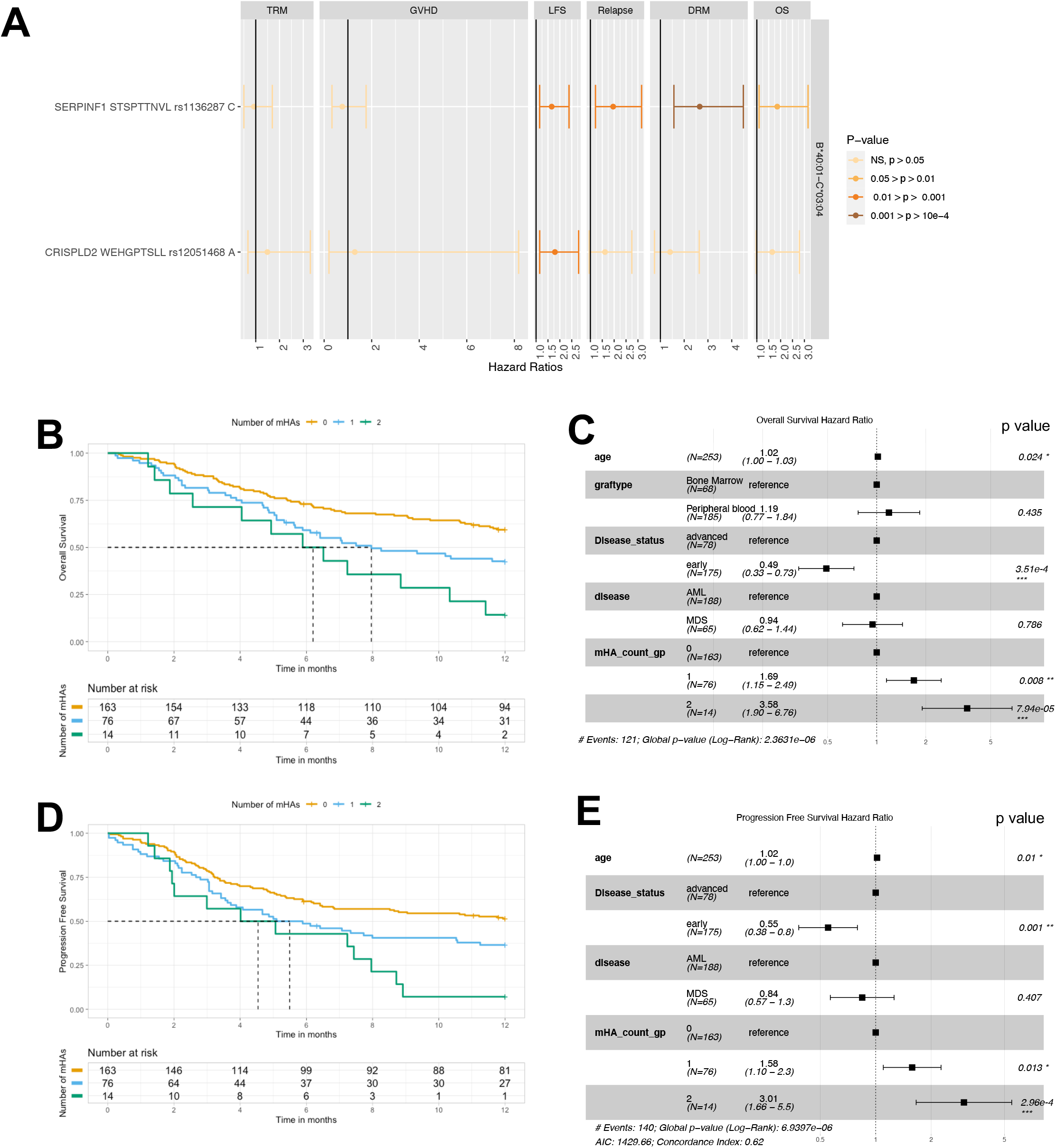
A multi-mHA analysis in an HLA haplotype shows a dose-response relationship between the number of mHAs present (0-2) and the hazard for various clinical outcomes. **(A)** Hazard ratios with 95% confidence interval from multi-mHA association analyses with 1-year post-transplant survival outcomes after meta-analysis of cohort 1 and 2. Each row represents the association of each mHA individually with the various outcomes (columns) in the patients with HLA haplotype B*40:01-C*03:04. **(B)** Kaplan-Meier curve showing differences in overall survival probability by mHA count groups. **(C)** Forest plot of the multi-mHA association model with overall survival. Each row shows the hazard ratio with 95% confidence interval of the variable. **(D)** Kaplan-Meier curve showing differences leukemia free survival probability by mHA count groups. **(E)** Forest plot of the multi-mHA association model with progression free survival. Each row shows the hazard ratio with 95% confidence interval of the corresponding variable.

## DISCUSSION

By implementing improved methods for mHA prediction in two large patient cohorts, we aimed to comprehensively explore the role of mHAs in alloHCT by analyzing whether (1) the number of predicted mHAs, or (2) individual mHAs are associated with clinical outcomes.

We found no correlation between the number of class I mHAs and the number of class II mHAs within DRPs. These two variables also had absent and weak correlations, respectively, with the number of mHA-mismatched SNPs within a DRP. These results suggest that the number of immunogenic mHAs predicted by our model was independent of the cumulative genetic differences between donors and recipients, and could explain why prior analyses did not find significant associations between genome-wide mismatching and clinical outcomes in unrelated donors.^13^ These data demonstrate that accurate characterization of the mHA landscape may require direct prediction and individual assessment of candidate peptides.

Recipients with a class I mHA count greater than the median had a 39% increased hazard of GvHD mortality compared to those with a number less than the median. While a previous study investigated the association of SNP-mismatching with GvHD, our results are the first to directly demonstrate that an increased number of predicted class I mHA peptides is associated with increased GvHD mortality following alloHCT.^13^

On the other hand, the number of class I mHAs was not significantly associated with outcomes indicative of GvL response – relapse, DRM, leukemia-free survival. There are several possible explanations for this. One possibility is that the class I GvL effect from engrafted T-cells is driven by immunodominant class I peptides rather than a cumulative effect of total class I mHAs. Previous studies have identified individual class I mHAs that have been found to elicit a strong anti-tumor response, including some that are being developed as therapeutic targets to bolster the GvL effect from transplants.^4^ Downregulation and loss of HLA molecules on tumor cells has been described as a mechanism for immune escape by tumor cells, and may also explain why we did not find associations between class I mHAs and increased GvL.^38^ It may also be possible that mHAs derived from SNP amino acid changes, as captured by our study, encompass only a small fraction of the landscape of peptides presented by tumor HLA molecules. Tumor cells also present antigens derived from overexpressed self-peptides, cancer/testis antigens, somatic variants, splicing variants, endogenous retroelements, alternative protein processing, and likely other sources as well as mHAs.^39-42^ It may be necessary to account for these components of the HLA-peptidome to uncover associations with GvL outcomes.

Survival analysis also showed that patients with a number of class II mHAs greater than the median had increased mortality compared to those with a number less than the median (KM P=0.03, CoxPH P=0.06). The consistently elevated hazard but the lack of statistical significance when adjusting for clinical covariates may point towards the need for more precise criteria for predicting class II mHAs or the need for a larger sample size. If this association is further validated, it may be due to recognition of class II mHAs by T-regulatory cells which diminish anti-tumor T-cell responses. But while mHA-specific T-regulatory cells have been isolated, the clinical significance of these is unknown.^43^

In an HLA-specific analysis, our study found three novel predicted class I mHAs and one class II mHA that are significantly associated with 1-year outcomes following alloHCT. DLRCKYISL (gene *GSTP1*, rs1695) was associated with increased 1-year GvHD mortality in recipients with HLA B*08:01. This SNP has long been under consideration around outcomes in both solid organ and stem cell transplant outcomes.^44^ The first study to investigate the role of this variant in HCT showed that a dominant model of rs1695 in donors was associated with increased risk for chronic GvHD incidence following matched-related donor alloHCT.^45^ However, this GvHD finding was not replicated at P<.05 in two large appropriately sized cohorts, including our DISCOVeRY-BMT study population.^46^ This failure to replicate when considering donors genetics highlights the complex interplay between donor and recipient genetics and the challenge in building data models to capture these relationships.

WEHGPTSLL (gene *CRISPLD2*, rs12051468) was associated with decreased overall survival in recipients with HLA-B*40:01. Though not statistically significant after Bonferroni correction, recipients with this mHA also had increased hazard for relapse and decreased overall survival. In recipients with HLA-C*03:04, STSPTTNVL (*SERPINF1*, rs1136287) was significantly associated with increased DRM. Patients with this mHA also had increased hazard for relapse and decreased overall survival that did not maintain statistical significance after multiple-testing correction. Our multi-mHA analysis within the HLA-B*40:01-C*03:04 haplotype showed that WEHGPTSLL and STSPTTNVL are additive contributors of mortality risk as there was a positive dose-dependent relationship between the number of mHAs present and the risks for decreased OS, increased DRM, and decreased LFS. These two mHAs, WEHGPTSLL and STSPTTNVL, had unexpected associations with 1-year outcomes. We expected mHAs to either reflect a GvH effect (i.e., increased GvHD mortality) or a GvL effect (i.e., decreased relapse and DRM; increased LFS). However, in both these cases, the associations reflect a diminished GvL effect. Similar findings have been previously reported – a study identified recipient allele mismatch associations (i.e., allele mismatches that have the potential to encode mHAs) that were associated with increased relapse in patients with HLA-A*02:01.^6^ It may be possible that these immunodominant mHAs compete with and limit HLA presentation of other immunogenic antigens and therefore suppress immune responses towards tumor cells, though there is no evidence to support this in this context. A similar mechanism has been described in the setting of cancer vaccination, where *Listeria monocytogenes* antigens have been theorized to suppress immunogenicity of target vaccine antigens.^47^

Our presented study has important limitations. The population was limited to DRP of EA ancestry and thus limiting generalizability of our findings. The frequencies of our mHA source SNP alleles and HLA alleles (Supplemental Table 3) vary by race and ethnicity; future analyses are planned and funded which will allow us to identify mHAs that may be more frequent across populations.^48,49^ Another limitation was that our model to predict mHAs relied on average gene expression from aggregate data across hundreds of samples.^27,28^ Further studies may require gene expression of individual patient tissue for more accurate mHA predictions. It may also be important to include predicted mHAs encoded by sources other than SNPs to assess the role of mHAs more comprehensively. It will also be important to include clinical outcomes beyond 1 year to investigate the long-term role of mHAs following alloHCT. Finally, it will be necessary to replicate our findings in other patient cohorts, and if they are consistent, they will need to be validated using *in vitro*/*vivo* studies.

## ACKNOWELDGMENTS

This study was supported by the National Heart, Lung, and Blood Institute (NHLBI) of the National Institutes of Health (NIH) (R01 HL102278; principal investigators, T.H. and L.E.S.-C.). This work was also supported by National Cancer Institute (NCI) grant P30CA016056, involving the use of the Roswell Park Cancer Institute Bioinformatics and Biostatistics shared resources. The authors appreciate funding from the American Society of Hematology HONORS Award and Medical Student Physician Scientist Award (OJ), and the Carolina Medical Student Research Program (OJ). The CIBMTR is supported primarily by Public Health Service U24CA076518 from the National Cancer Institute (NCI), the National Heart, Lung and Blood Institute (NHLBI) and the National Institute of Allergy and Infectious Diseases (NIAID); U24HL138660 and U24HL157560 from NHLBI and NCI; U24CA233032 from the NCI; OT3HL147741 and U01HL128568 from the NHLBI; HHSH250201700005C, HHSH250201700006C, and HHSH250201700007C from the Health Resources and Services Administration (HRSA); and N00014-20-1-2832 and N00014-21-1-2954 from the Office of Naval Research. The views expressed in this article do not reflect the official policy or position of the National Institute of Health, the Department of the Navy, the Department of Defense, Health Resources and Services Administration (HRSA) or any other agency of the U.S. Government.

## AUTHORSHIP

O.J., H.T, L.E.S.-C designed and performed research. O.J. and H.T. analyzed and interpreted data and performed statistical analysis. O.J. wrote the manuscript. K.O., P.A., and B.V. interpreted data and provided feedback on research design and analysis. S.V. created computational pipelines. Q.Z. provided quality control and data randomization. Y.W. provided quality control. C.A.H., L.P., and X.S. performed genotyping interpretation of data. G.B., A.W. performed quality control and coding. M.C.P., P.L.M., S.R.S., T.H. acquired, organized and adjudicated clinical data. All authors revised the manuscript, contributed critically important intellectual content, and approved the final version of the manuscript.

---

**Supplemental Figure 1.**
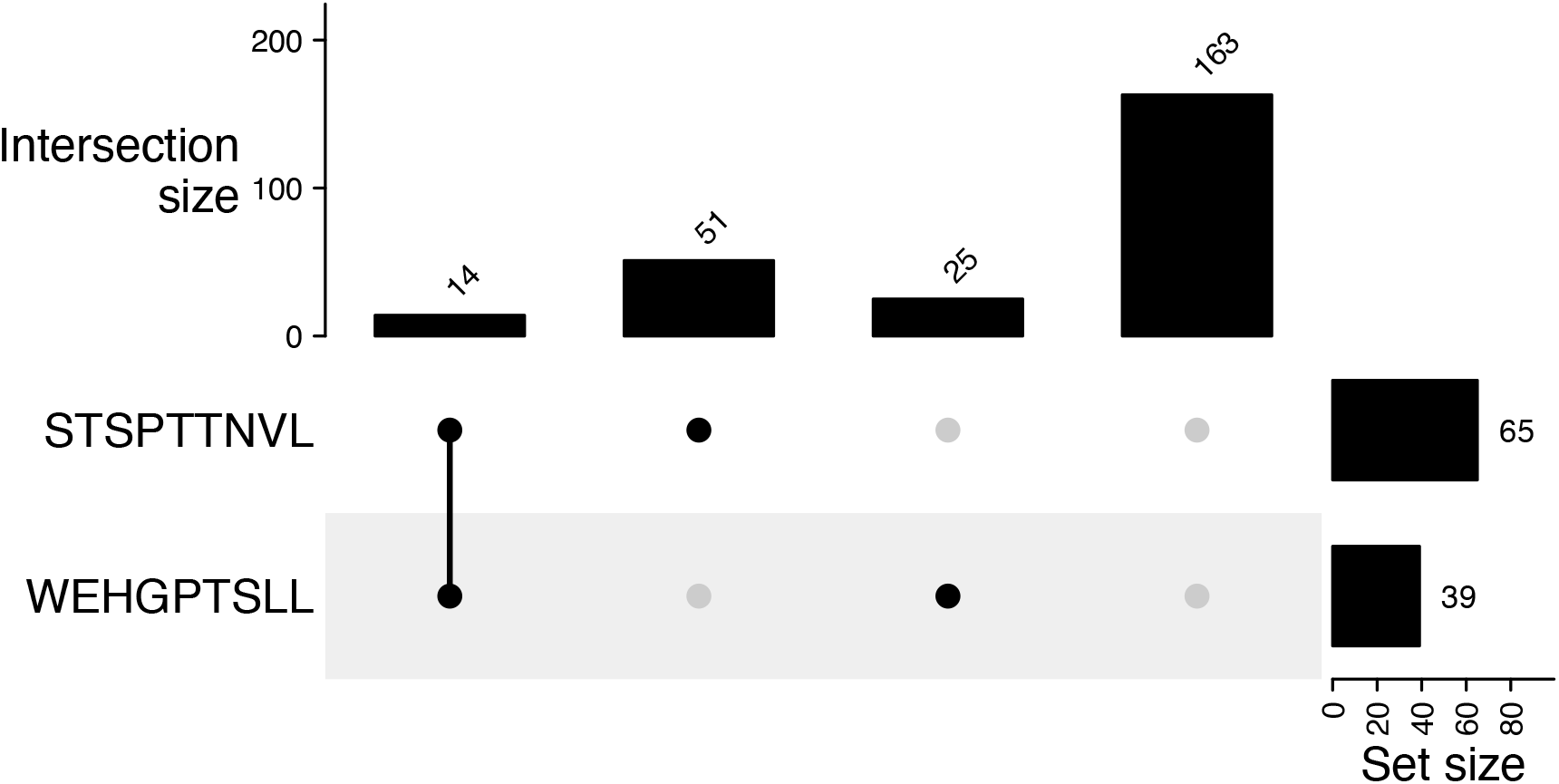
The number of donor recipient pairs with haplotype HLA B*40:01-C*03:04 that contain various combinations of the mHAs WEHGPTSLL and STSPTTNVL.

**Supplemental Figure 2.**
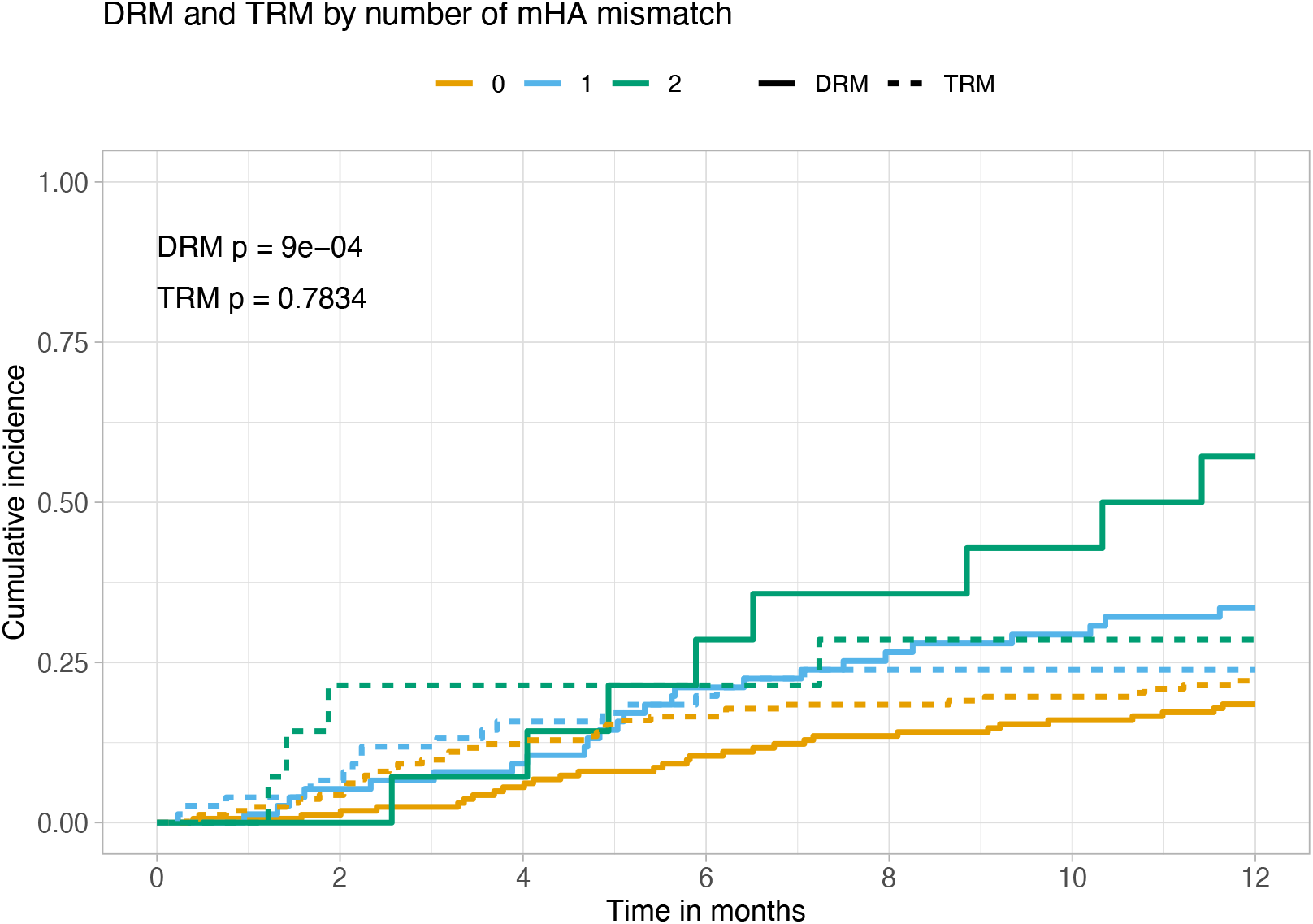
Cumulative events of disease related mortality (DRM) and transplant related mortality (TRM) stratified by the number of mHAs, WEHGPTSLL and STSPTTNVL, in DRPs with HLA B*40:01-C*03:04.

**Supplemental Table 1.** Model covariates used in survival models. This table displays representative coefficients, hazard ratios, and p values from class I quantile survival analyses. Values of NA indicate a covariate was not used in models for that given outcome.

|  | OS |  |  | LFS |  |  | DRM |  |  | Relapse |  |  | TRM |  |  | GvHD Mortality |  |  |
| --- | --- | --- | --- | --- | --- | --- | --- | --- | --- | --- | --- | --- | --- | --- | --- | --- | --- | --- |
| Covariates | HR | 95% CI | P value | HR | 95% CI | P value | HR | 95% CI | P value | HR | 95% CI | P value | HR | 95% CI | P value | HR | 95% CI | P value |
| Recipient Age | 1.014 | 1.01 - 1.019 | 5.68 x 10 | 1.012 | 1.008 - 1.017 | 3.89E-08 | 1.01 | 1.0005 - 1.019 | 0.0395 | 1.006 | 1.0007 - 1.01 | 0.0265 | 1.016 | 1.009 - 1.023 | 6.31E-06 | 1.018 | 1.008 - 1.028 | 0.00033 |
| Donor Age | NA | NA | NA | NA | NA | NA | NA | NA | NA | NA | NA | NA | NA | NA | NA | 1.014 | 1.00 - 1.028 | 0.05 |
| MDS | 0.809 | 0.689 - 0.95 | 0.0098 | 0.808 | 0.688 - 0.949 | 0.0092 | 0.46 | 0.358 - 0.59 | 9.67E-10 | 0.529 | 0.431 - 0.649 | 1.05E-09 | 1.471 | 1.19 - 1.82 | 0.00033 | 1.928 | 1.44 - 2.57 | 8.50E-06 |
| Early disease | 0.542 | 0.474- 0.621 | 6.72E-19 | 0.535 | 0.467 - 0.612 | 8.97E-20 | 0.359 | 0.3 - 0.43 | 4.88E-29 | 0.44 | 0.377 - 0.514 | 4.68E-25 | NA | NA | NA | NA | NA | NA |
| Peripheral blood graft | 0.772 | 0.668 - 0.893 | 0.00048 | NA | NA | NA | NA | NA | NA | NA | NA | NA | 0.684 | 0.549 - 0.852 | 0.00071 | NA | NA | NA |
| Obese BMI | NA | NA | NA | NA | NA | NA | NA | NA | NA | NA | NA | NA | 1.28 | 1.0028 - 1.63 | 0.047 | 1.36 | 0.95 - 1.94 | 0.089 |
| Overweight BMI | NA | NA | NA | NA | NA | NA | NA | NA | NA | NA | NA | NA | 1.047 | 0.81 - 1.35 | 0.72 | 1.13 | 0.78 - 1.63 | 0.52 |

**Supplemental Table 2.** Sub-distributional hazard models for relapse, DRM, and TRM in DRPs with haplotype HLA B*40:01-C*03:04. Included as a variable is the number of mHAs, WEHGPTSLL and STSPTTNVL, a DRP has. “mHA count=2” signifies a DRP has both of these mHAs, while “mHA count=1” signifies a DRP has either one of these mHAs. We observe a dose-response relationship between the mHA count and increased hazards for relapse, DRM, and TRM.

| Characteristic | Relapse |  |  | DRM |  |  | TRM |  |  |
| --- | --- | --- | --- | --- | --- | --- | --- | --- | --- |
|  | HR<sup>†</sup> | 95% CI<sup>†</sup> | p-value | HR<sup>†</sup> | 95% CI<sup>†</sup> | p-value | HR<sup>†</sup> | 95% CI<sup>†</sup> | p-value |
| Recipient age | 1.01 | 0.99, 1.03 | 0.2 | 1.01 | 0.99, 1.04 | 0.2 | 1.02 | 1.00, 1.04 | **0.030** |
| Advanced disease | 1.67 | 1.04, 2.68 | **0.035** | 2.18 | 1.27, 3.74 | **0.005** |  |  |  |
| AML | 1.55 | 0.90, 2.67 | 0.11 | 1.51 | 0.82, 2.78 | 0.2 | 0.75 | 0.43, 1.31 | 0.3 |
| mHA count = 1 | 1.71 | 1.07, 2.74 | **0.025** | 2.05 | 1.21, 3.47 | **0.008** | 1.08 | 0.61, 1.92 | 0.8 |
| mHA count = 2 | 2.85 | 1.43, 5.68 | **0.003** | 3.65 | 1.57, 8.51 | **0.003** | 1.32 | 0.42, 4.15 | 0.6 |
| Obese |  |  |  |  |  |  | 0.59 | 0.31, 1.14 | 0.12 |
| Overweight |  |  |  |  |  |  | 0.51 | 0.26, 0.97 | **0.039** |
| Peripheral blood |  |  |  |  |  |  | 0.68 | 0.38, 1.21 | 0.2 |

<sup>†</sup> HR = Hazard Ratio, CI = Confidence Interval

**Supplemental Table 3.** Frequencies of mHA source-SNP alleles and HLA allele/haplotype among European Americans and African Americans.

| mHA<br>SNP allele<br>HLA allele | European Americans |  | African Americans |  |
| --- | --- | --- | --- | --- |
|  | SNP allele<br>frequency | HLA allele/haplotype<br>frequency | SNP allele<br>frequency | HLA allele/haplotype<br>frequency |
| DLRCKYISL<br>rs1695, ref, G<br>HLA B*08:01 | 0.6673 | 0.12525 | 0.5523 | 0.03838 |
| WEHGPTSLL<br>rs12051468, ref, A<br>HLA B*40:01 | 0.5651 | 0.05643 | 0.6044 | 0.01328 |
| STSPTTNVL<br>rs1136287, ref, C<br>HLA C*03:04 | 0.3539 | 0.08215 | 0.1627 | 0.0533 |
| YQEIAAIPSAGRERQ<br>rs11200385, alt, A<br>HLA DRB1*15:01 | 0.2279 | 0.14441 | 0.3383 | 0.02931 |
| HLA B*40:01-HLA C*03:04 | NA | 0.05538 | NA | 0.01226 |

## Notes

### Competing Interest Statement

The authors have declared no competing interest.

